# Unlocking efficiency: Native circular RNA surpass linear isoforms in RNase P activity

**DOI:** 10.1101/2025.06.30.662392

**Authors:** A.K. Chawla, A.M. Kietrys

## Abstract

RNase P, one of the earliest enzymes identified with RNA as its catalytic component, has been extensively studied across all three domains of life. In this research, we unveil circular isoforms of RNase P RNA within bacterial, fungal, and human cell lines. Comparing the bacterial variant, circM1, with its linear counterpart under diverse conditions revealed its enhanced temperature resistance and superior tolerance to Mn^2+^. Moreover, our findings suggest distinct protein associations for both isoforms in the presence of FBS. The human counterpart, circH1, was proved to be active *in cellulo*.

## Introduction

One of the only two known universal ribozymes^1,2^, RNase P has been proved to catalyze the maturation of precursor-tRNA (pre-tRNA) in a Mg^2+^ dependent manner^3^ (Fig. 1). Across all domains, it consists of one non-coding catalytic RNA and varying number of proteins. In human cells, it has further been shown to play a crucial role in RNA polymerase III transcription through the formation of initiation complexes^4^. The structure and catalytic activity of the bacterial RNase P have been studied extensively both *in vitro* and *in cellulo*^5–7^. The catalytic component, M1 RNA, is catalytically active independent of the RNase P protein *in vitro*. The human counterpart, H1 RNA was found to have little activity *in vitro* in the absence of its associate proteins in a high Mg^2+^ concentration (400mM) over 23 hours^8^. Yeast RNase P RNA, Rpr1 hasn’t yet been reported to be active independent of its nine cognate proteins.

**Figure 1.**
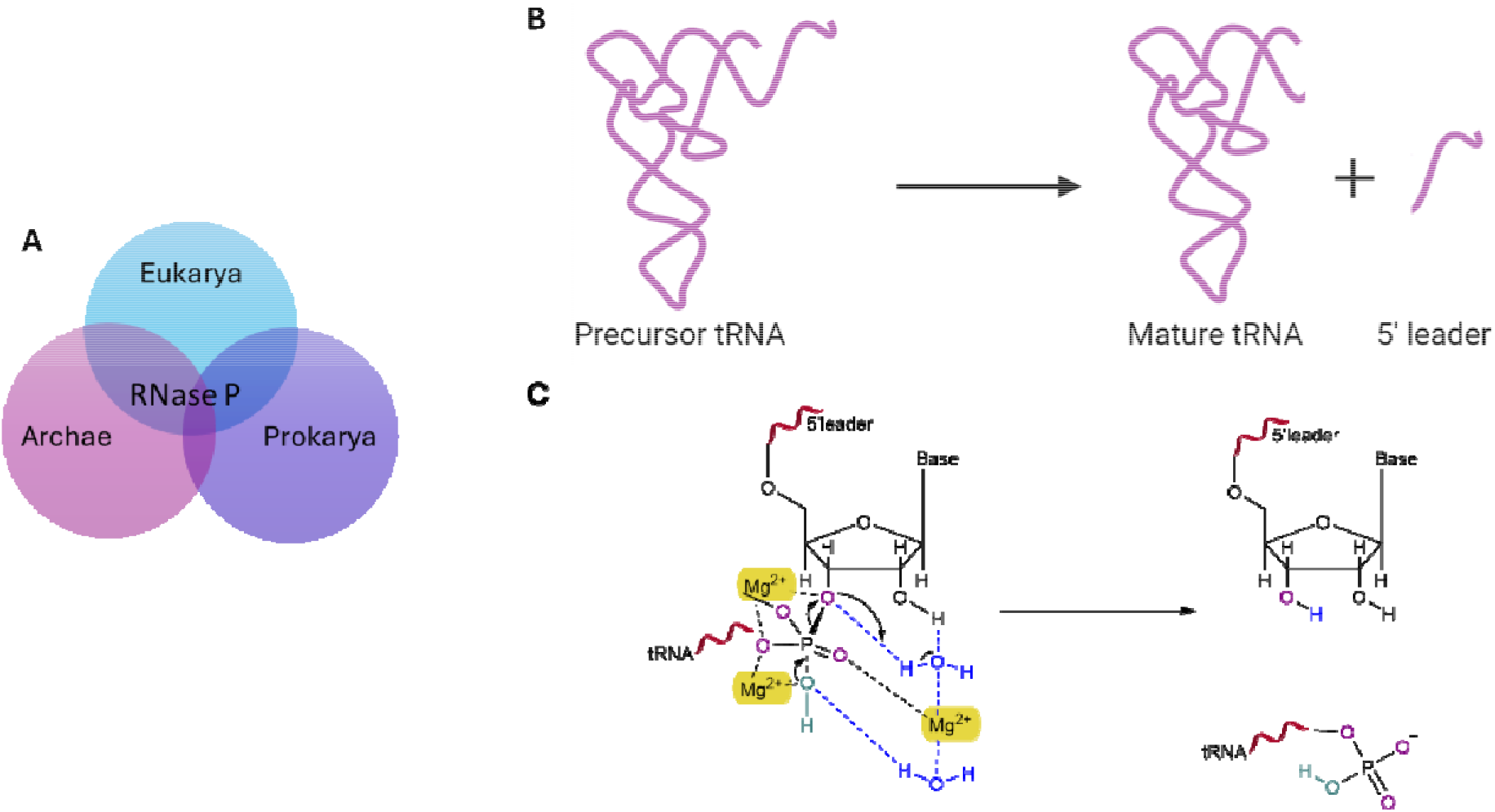
RNase P mediates maturation of pre-tRNA. (A) RNase P is a universal ribozyme existing in all three domains of life-prokarya, archae and eukarya. (B) A 5’ leader is cleaved off precursor tRNA, an essential step i tRNA maturation. (C) RNase P enzymatic activity is Mg^2+^ dependent. Three Mg^2+^ hold the pre-tRNA in proximity of a water molecule to facilitate hydrolysis at the cleavage site.

Besides ribozymes, circular RNAs (circRNAs) are another class of non-coding RNA. However, circRNAs have not been as extensively studied due to their fairly recent discovery^9^. They have been shown to have longer half-lives and higher resistance against exonucleases when compared to their linear counterparts^10,11^. Studies have found circRNAs to play crucial roles in gene regulation^12^, translation^13^, microRNA sponging^14^, signaling^15^ and disease progression^16,17^. CircRNAs derived from the H1 RNA gene, RPPH1, have been implicated in breast cancer^18^, colorectal cancer^19^ and cervical cancer^20^. While circular ribozymes have been synthesized *in cellulo* through the permuted intron-exon (PIE) system^21^, no natively occurring circular ribozymes have been reported up until now.

In this study, we have reported the first natively occurring circular ribozyme. We discovered circular variants of RNase P RNAs in bacterial, fungal and human cell lines (Scheme 1). We further investigated the *in vitro* activity of the bacterial variant, circM1 and *in cellulo* activity of the human variant, circH1.

**Scheme 1.**
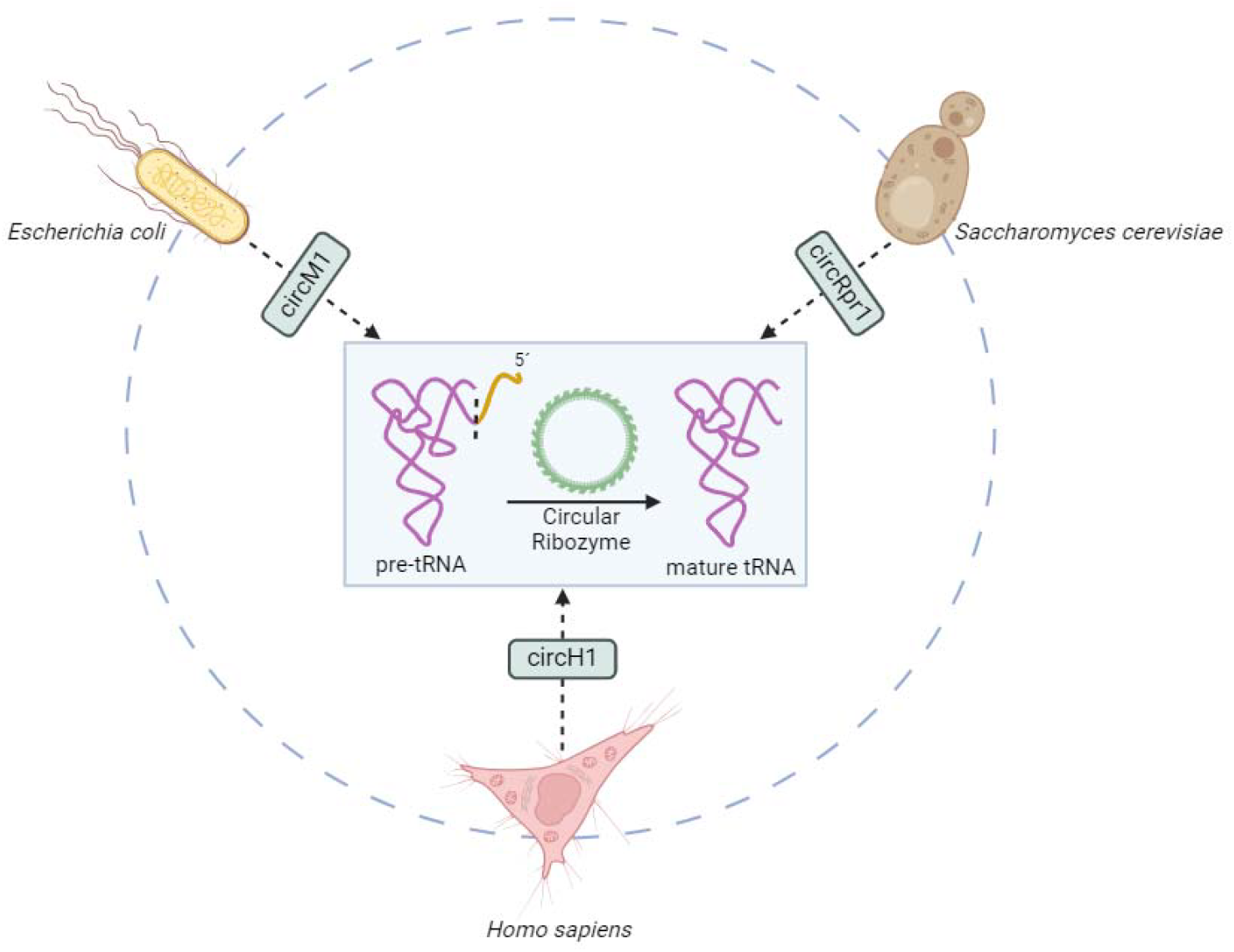
Circular RNase P RNA was found in three different species, namely *E. coli, S. cerevisiae* and *H. sapiens*

### Circular variants of RNase P RNA were found in bacterial, fungal and human cell lines

In the search of circRNA present abundantly in human brain, we found hsa_circ_0000514 which is derived from the RPPH1 gene locus^22,23^. On further examination using publicly available databases^24,25^, we found 27 reported circRNA being generated from this locus. We then set to find if there exists a circular variant of linear transcript, H1. Using divergent primers, we were successful in detecting circH1 in HeLa cells. To verify the circular nature of the detected transcript, RNase R treatment was performed on total HeLa RNA, the results of which proved the existence of circH1 (Fig. 2B). Secondary structures predicted using RNAfold^26^ (Fig. S1) were identical for the two isoforms, which further increased our curiosity in the function of this circular RNA. Since RNase P is a universal ribozyme, we set forth to investigate if the circular isoform is also universal across all domains. Using the same experimental strategy as described above, circM1 and circRpr1 were confirmed to be present in *Escherichia coli* (*E. coli*) and in *Saccharomyces cerevisiae* (*S. cerevisiae*), respectively (Fig. 2C, D). Next, we investigated the sequence similarity of the linear and circular counterparts to identify conserved regions between the two isoforms. Divergent primers were utilized to amplify the entirety of the circRNA sequences using Polymerase Chain Reaction (PCR). The PCR products were analyzed using Sanger sequencing. The alignment was in excellent agreement with the known linear sequences. CircH1 aligned with 1 mismatch, circM1 with 4 mismatches and circRpr1 with multiple mismatches not aligning on the two strands indicating possible RNA modifications.

**Figure 2.**
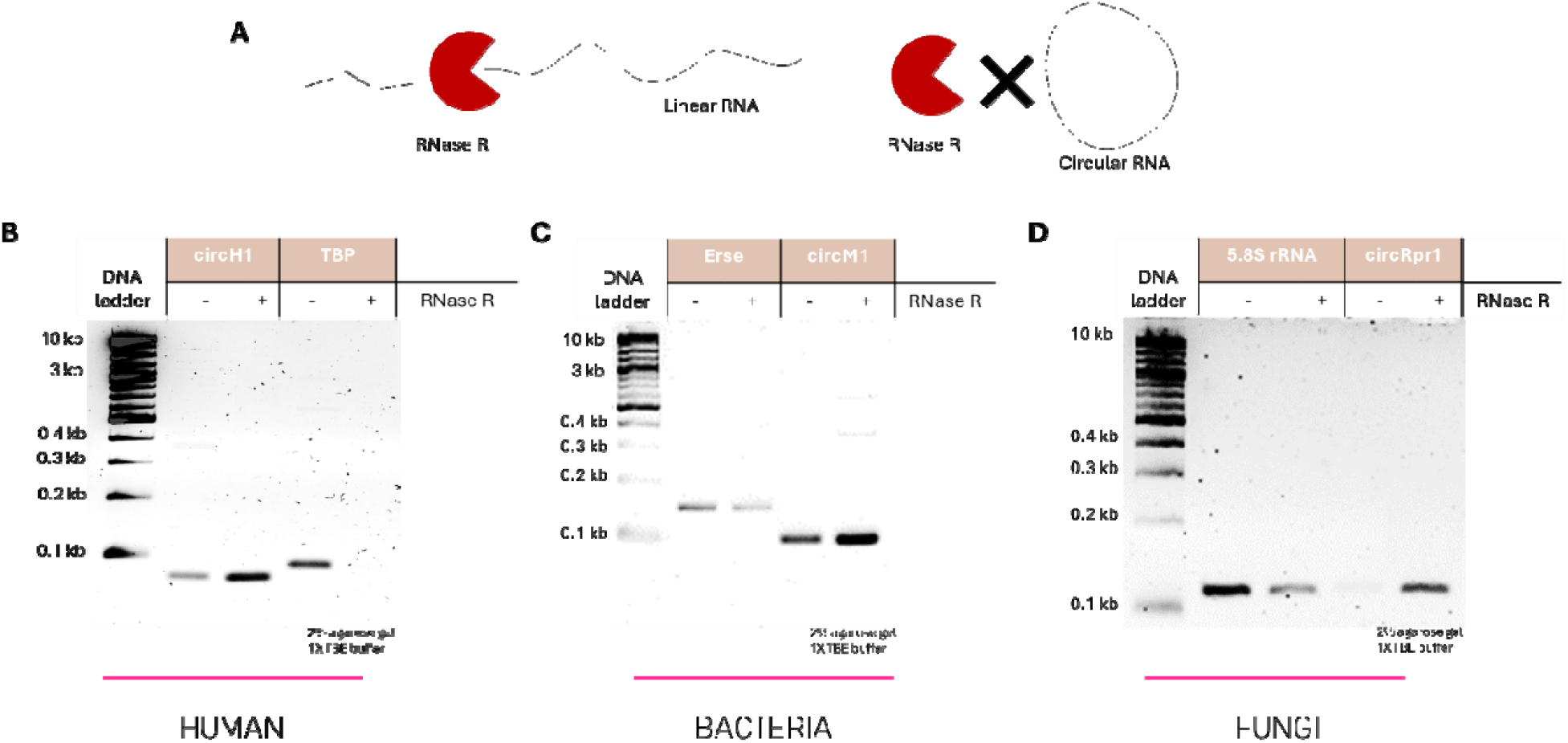
RNase P circRNA were verified in three species using RNase R treatment. (A) RNase R is a 3’ exonuclease capable of cleaving linear RNA but not circular RNA. The presence of circH1 (B), circM1 (C) and circRpr1 (D) was verified using RNase R treated total RNA extracted from H. sapiens, E. coli and S. cerevisiae respectively. TBP (B), Erse (C) and 5.8S rRNA (D) were used as positive controls. (+) denotes total RNA treated with RNase R and (−) denotes untreated total RNA. Following RNase R treatment, cDNA synthesis was performed followed by PCR using specific primers. PCR products were analyzed on 2% agarose gels.

### CircM1 outperforms linM1 under numerous conditions

Since circRNAs have been reported to have higher stability and structural rigidity in comparison to linear RNAs, we next explored the comparative activity of the discovered linear and circular ribozymes. Since catalytic activity of linM1 has been extensively studied, we based our in vitro comparative studies on linM1 versus circM1. The reactions were performed on an extensively studied model pre-tRNA substrate, pATSerUG (Fig. 3B). The two isoforms exhibited similar activities at the previously reported optimized conditions for linM1 (Fig. 3A). While both linM1 and circM1 perform best at 37°C and lose activity at extremes, circM1 has a higher resistance to an increase in temperature from the optimal of 37°C (Fig. 3C). This could be explained by a lower entropy advantage for circM1 upon unfolding due to its closed loop structure. Since the catalytic mechanism of RNase P is Mg^2+^ dependent, we next investigated if the two M1 isoforms correspond differently along a Mg^2+^ concentration gradient. We found both isoforms to have an increase in activity with an increasing [Mg^2+^] concentration. While both linM1 and circM1 have comparable performances at [Mg^2+^] up to 40 mM, concentrations tested above this bench point showed that circM1 responds more to an increase in the [Mg^2+^] as compared to linM1 (Fig. 3D). Previous studies have shown that substitution of Mg^2+^ by Mn^2+^ can have adverse effects to the catalytic activity of linM1^27^. We then varied the [Mg^2+^]/[Mg^2+^+Mn^2+^] in the reaction. As expected, both linM1 and circM1 perform better under low [Mn^2+^]/high [Mg^2+^] conditions. Interestingly, circM1 is more tolerant to Mn^2+^ than linM1 as is evident at [Mg^2+^]/[Mg^2+^+Mn^2+^] of 0.5 (Fig. 3E). This tolerance could be a result of circM1’s capability to selectively exclude the Mn^2+^ ions and more efficiently incorporate the available Mg^2+^ ions for the catalytic cleavage. Since circRNAs have been shown to have longer half-lives, we next investigated the time-dependency of the activity of the two M1 isoforms. Contrary to our expectations, circM1 had a higher activity than linM1 in the initial time points. LinM1 took more than 60 minutes to catch up with circM1. We didn’t see a difference in activities over longer periods (Fig. 3F). This can be explained by the high activity rate of the reaction; by the time the catalytic RNAs start degrading, the percent cleavage plateaus. Furthermore, the turnover rate of circM1 could be higher than linM1 causing the significant difference during the initial time points. pH has also been shown to affect the catalytic activity of linM1, with a pH 7.2 corresponding to the maximum activity^28^. We observed a similar trend for circM1, with no apparent differences between the pH profiles of the two M1 isoforms (Fig. S5).

**Figure 3.**
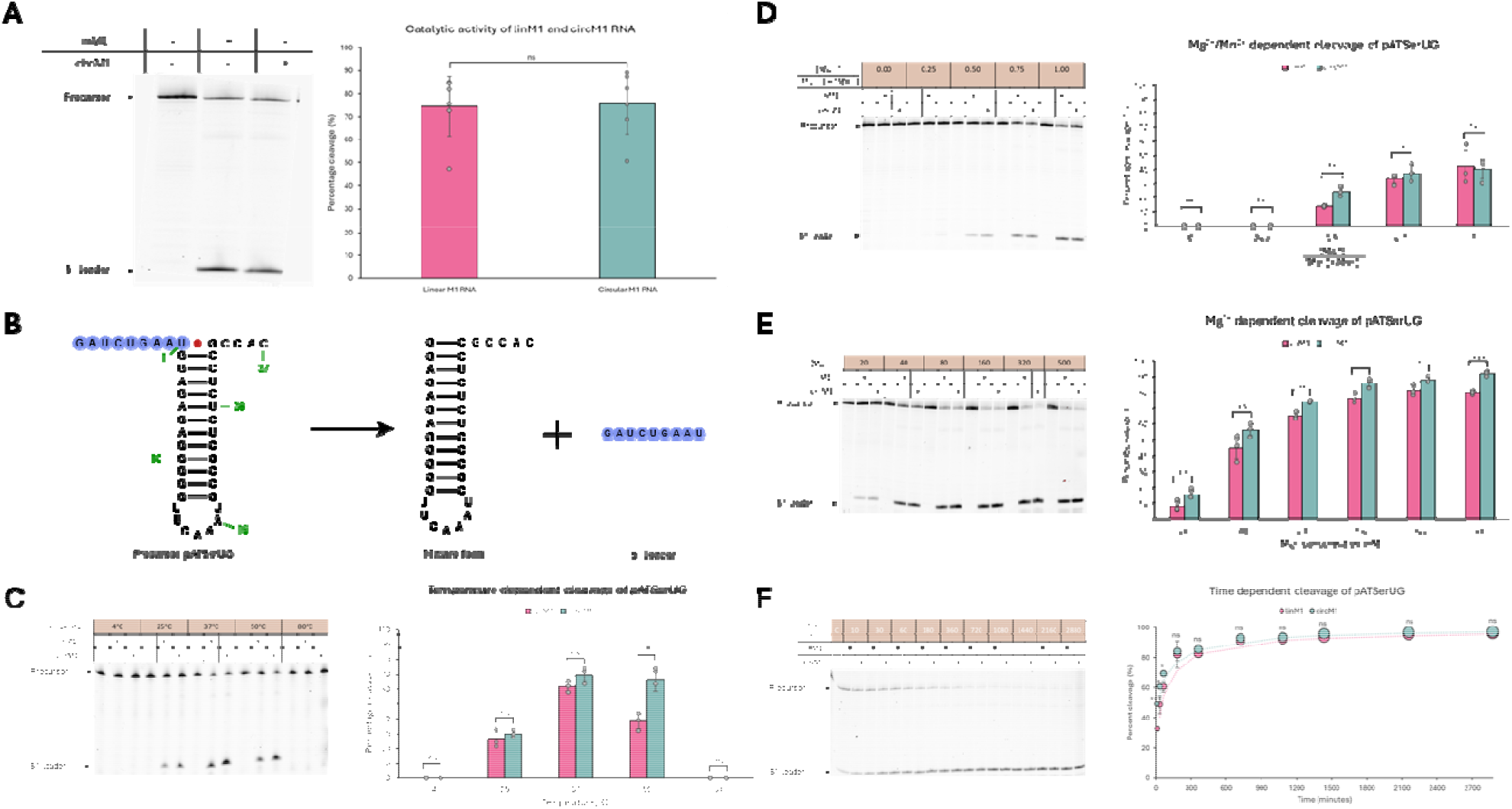
LinM1 and circM1 enzymatic activities were compared under different conditions. (A) Buffer (40mM MgCl_2_, 50mM Tris HCl pH 7.2, 5% PEG8000, 100mM NH_4_Cl), 0.24µM M1 RNA, 0.05µM 5’Cy5-pATSerUG. M1 RNA wa incubated in buffer for 30 minutes at 37°C followed by addition of pATSerUG pre-warmed to 37°C. Reaction (2µL) was incubated at 37°C for 40 minutes, quenched with 8µL of 2X Orange G-formamide and analyzed on a 20% denaturing polyacrylamide gel. (B) pATSerUG is a model pre-tRNA with a 9 nucleotide long leader sequence. (C) Reaction temperatures of 4°C, 25°C, 37°C, 50°C and 80°C were tested (D) [Mg^2+^]/[Mg^2+^]+[Mn^2+^] of 0.00, 0.25, 0.50.0.75 and 1.00 were tested (E) Mg^2+^ of 20 mM, 40mM, 80mM, 160mM, 320mM and 500mM were tested. (F) Reaction time periods of 10min, 30 min, 60 min, 3 hrs, 6 hrs, 12 hrs, 18 hrs, 24 hrs, 36 hrs, 48 hrs were tested. For all quantitative analyses, n=3. Student’s t-test were performed to calculate p-values. p>0.05, ns; p≤0.05, ^*^; p≤0.01,^**^; p≤0.001,^***^.

### M1 isoforms behave differently in FBS

Since circRNAs have been shown to be more resistant to exonuclease degradation than their linear counterparts, we next wanted to evaluate the activity of linM1 and circM1 in a nuclease rich environment. The pre-tRNA maturation reaction was performed in 1% FBS. Our initial hypothesis was that circM1 should perform better than linM1 due to its enhanced stability. Interestingly, we found that circM1 and linM1 behave differently. In the absence of FBS, only the canonical 9-mer product is observed. However, in the presence of 1% FBS, we see two products, the 9-mer leader (Fragment 1) and a shorter (<9 nt) fragment (Fragment 2). Compared to circM1, linM1 promotes the formation of the Fragment 1 more. On the other hand, Fragment 2 production is enhanced in the presence of circM1 when compared to linM1 (Fig. 4)

**Figure 4.**
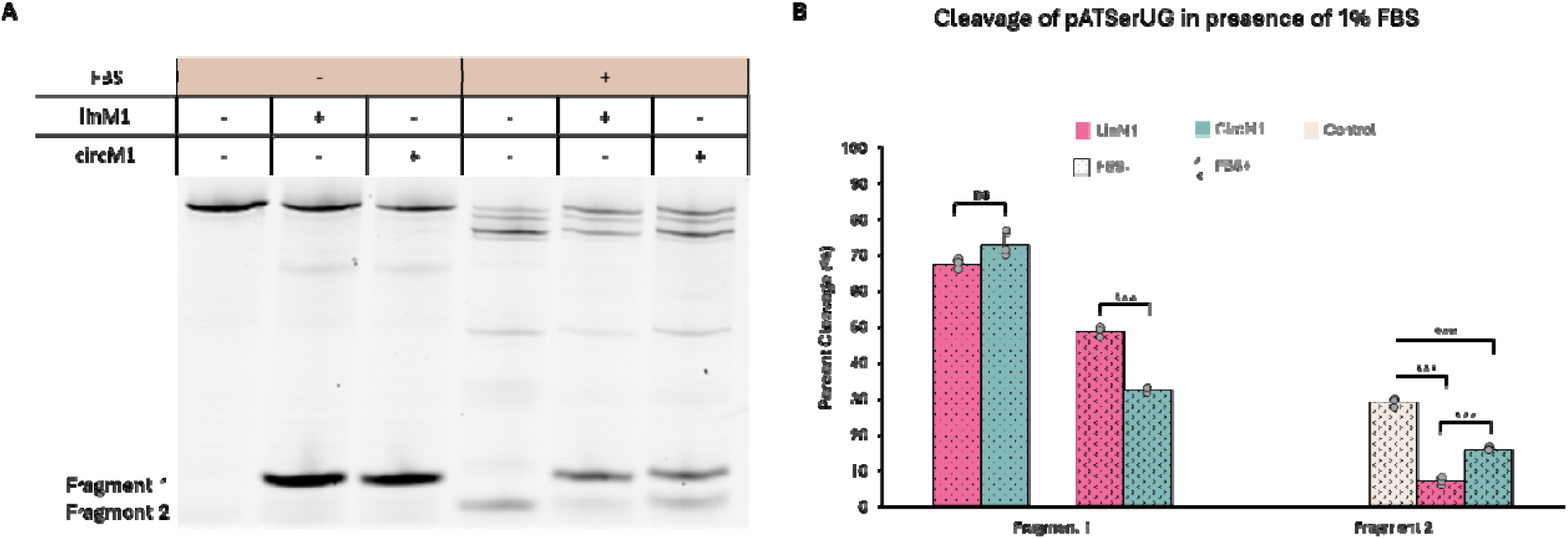
LinM1 and circM1 activities in presence of 1% FBS show difference in cleavage site preference. (A) Buffer (40mM Mg^2+^, 50mM Tris HCl pH 7.2, 5% PEG8000, 100mM NH_4_Cl), 0.24µM M1 RNA, 0.05µM 5’Cy5-pATSerUG, 1% FBS. M1 RNA was incubated in buffer for 30 minutes at 37°C followed by addition of FBS, and pATSerUG pre-warmed to 37°C. (B) Quantitative analysis shows significant difference in cleavage site preference of the two M1 isoforms in 1% FBS. For quantitative analysis, n=3. Student’s t-test were performed to calculate p-values. p>0.05, ns; p≤0.05, *; p≤0.01,**; p≤0.001, ***.

1. suppression of circM1 in FBS or 2. different protein, nucleic acid, small molecule or ion associations of the two isoforms.

### RNase P circRNA levels are equivalent to linear RNA levels in eukaryotes

Global circRNA levels have been shown to be significantly lower than linear RNA levels. In particular, most genes have much higher linear isoform levels compared to their linear counterparts with a tissue-specific exceptions such *Rims2*^*29*^. Using RT-qPCR in conjunction with 2^−ΔΔCt^ method for data analysis and TBP, 5srRNA and Erse were used as housekeeping genes for HeLa, *S. cerevisiae* and *E. coli* cells. Given the superior stability of circRNA, our initial hypothesis was that these circRNA are relics of the RNA world. To our astonishment, RNase P circRNA levels relative to the linear isoform was extremely low in bacteria less than 1%. However, the circRNA levels were equivalent (~49% in HeLa) or higher (~62% in *S. cerevisiae*) compared to the RNase P linear RNA levels (Fig. 5c). This finding suggests that these catalytic circRNA are not relics of the past but recently developed advanced mechanisms as evident by their high levels in eukaryotic organisms.

**Figure 5.**
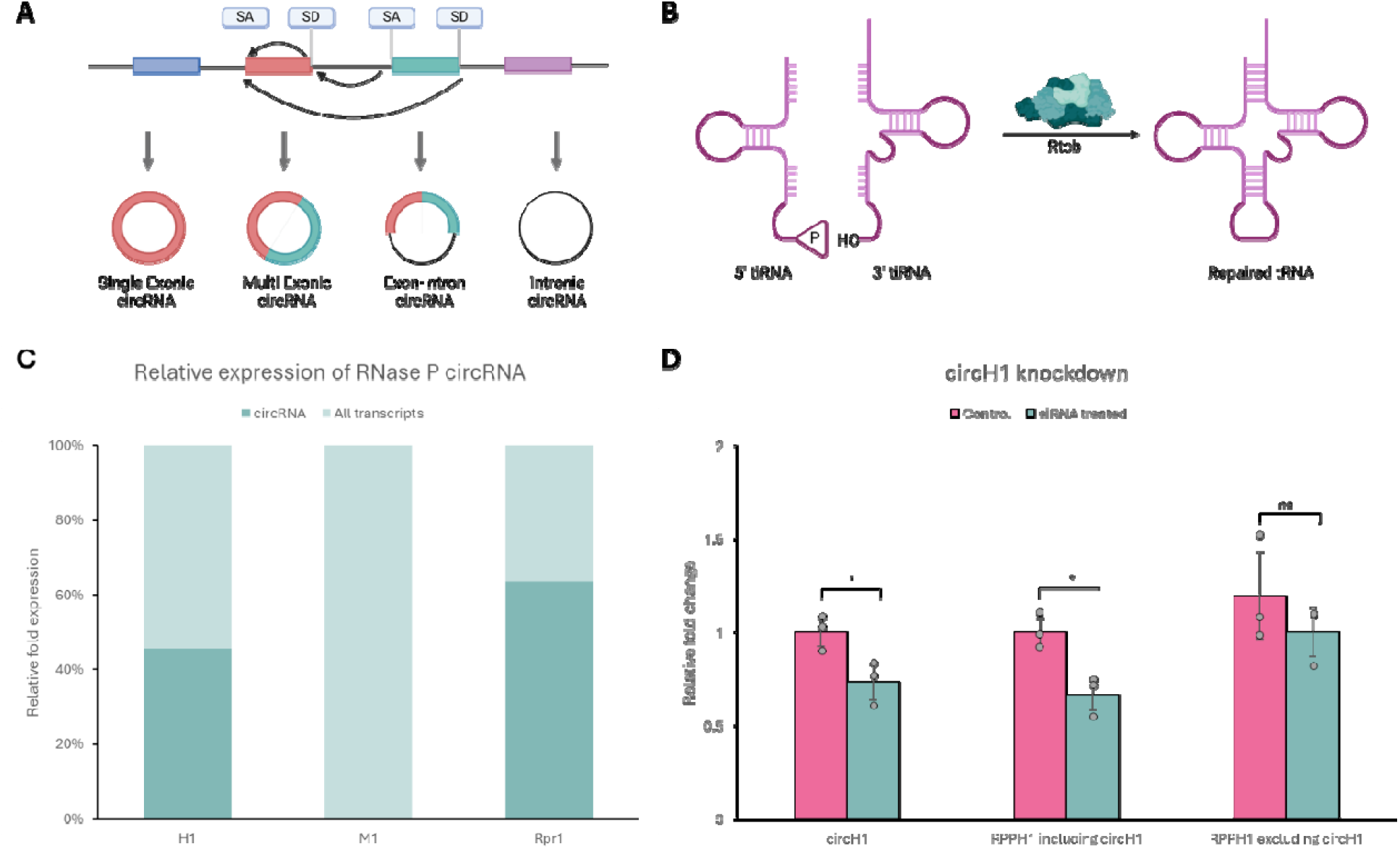
RNase P circRNA are highly expressed in eukaryotes (A) CircRNA are generally produced throug backsplicing; Splice donor (SD) attacks splice donor (SD) leading to the formation of different types of circRNA (B) Rtcb1 produces tRNA intronic circular RNAs through ligation of 5’-hydroxyl and 2’, 3’-cyclic phosphate at the 3’ end.(C) CircH1, circM1 and circRpr1 levels were compared to all H1, M1 and Rpr1 transcripts using RT-qPCR in Hela, S. *cerevisiae* and DHα E. *coli* cell lines using TBP, 5s rRNA and Erse genes respectively. (D) Quantitative analysis confirms specific knockdown of circH1 using BSJ-targeted siRNA knockdown. For quantitative analysis, n=3. Student’s t-test were performed to calculate p-values. p>0.05, ns; p≤0.05, *; p≤0.01,**; p≤0.001, ***.

**Figure 5a.**
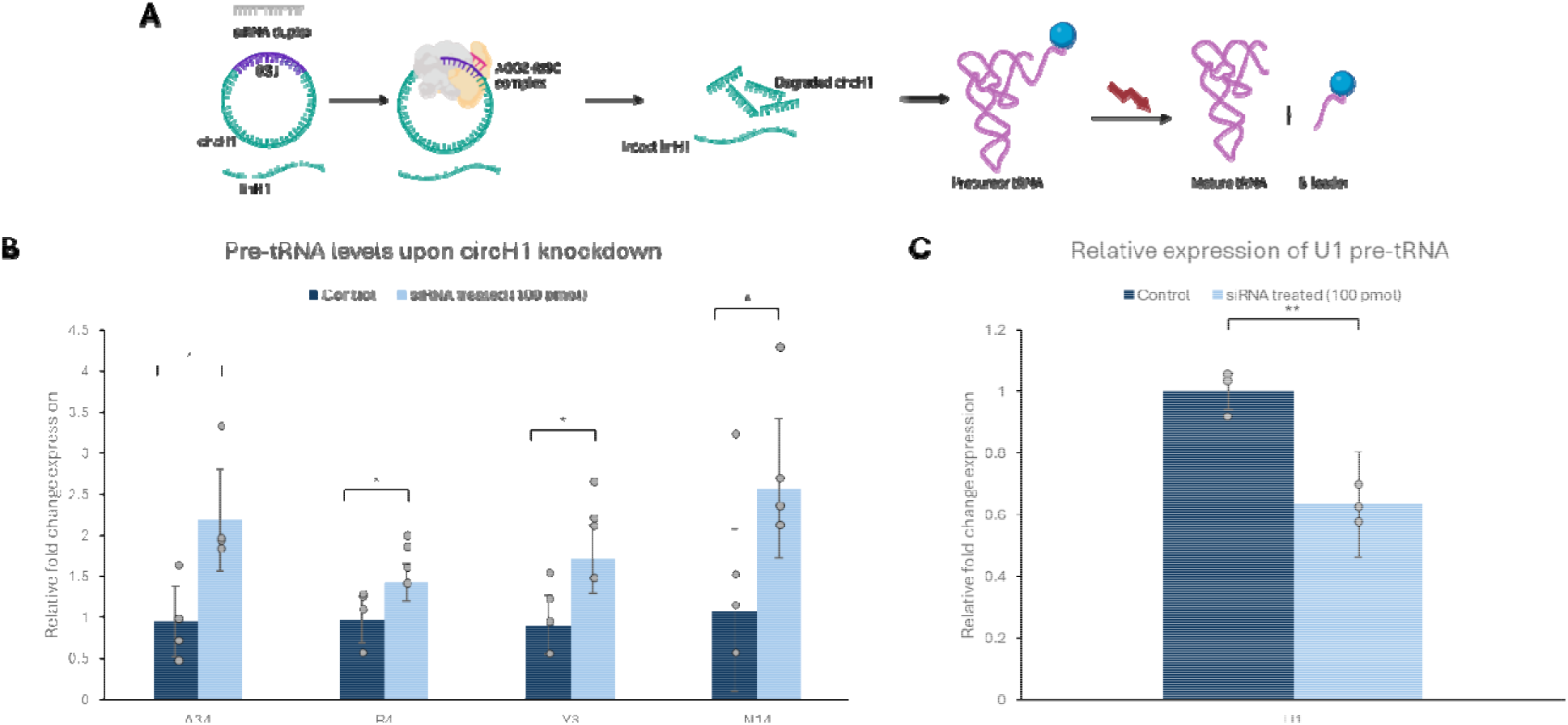
CircH1 is involved in pre-tRNA maturation. (A) siRNA was designed to specifically target BSJ of circH1 leaving linH1 intact. A decreased level of circH1 corresponds to increased levels of pre-tRNA levels—a direct evidence for circH1 involvement in pre-tRNA maturation (B) Quantitative analysis shows significant increase in pre-tRNA levels upon circH1 knockdown. (C) circH1 knockdown leads to significant selenocysteine pre-tRN downregulation. For quantitative analysis, n=4. Student’s t-test were performed to calculate p-values. p>0.05, ns; p≤0.05, *; p≤0.01,**; p≤0.001, ***.

We next evaluated if circRNA levels can affect linear RNA levels and if the production of these two isoforms is interdependent. Since siRNA knockdown in human cell lines is well documented to knockdown circRNA, we chose to perform circRNA downregulation in HeLa cells. SiRNA was designed to target the backsplice junction of circH1. We first evaluated the efficiency of siRNA knockdown and through RT-qPCR. We observed a significant knockdown of circH1 at siRNA concentration of 100nM. However, linH1 levels remained constant (Fig. 5d). Since the circRNA sequences in this study have a high similarity to their linear counterparts and the fact that the H1 gene does not contain any introns, backsplicing doesn’t seem like a plausible mechanism for circH1 production (Fig. 5a). This finding further concretes the unlikeliness of backsplicing being the mechanism for circH1 production and that circH1 production is post-transcriptional. Rtcb is a highly conserved protein shown to ligate tRNAs with a phosphorylated 3′-end and the other with a hydroxylated 5′-end leading to the formation of tRNA intronic circular (tric) RNAs through a backsplicing-independent mechanism^30^. Rtcb-like proteins are known to be present in archaea, bacteria, algae and animals but absent in fungi and plants^31^ (Fig. 5b). We believe that circH1 production happens through a similar mechanism, however, this needs to be validated by evaluating the protein interactome of circH1 and studying the direct effects of downregulation of any potential protein candidates.

Linear H1 has been shown to localize throughout the nucleus and the cytoplasm^32^. The localization of the RNA heavily influences the function it is involved in^33^. One established method to determine the localization of RNA molecules in a cell is FISH^34^. However, to obtain observable fluorescence, multiple probes are used to target the different sequences of the RNA and “lighting” it up enough to visualized using fluorescence microscopy^35^. Due to the high sequence similarity of linear and circH1, we were limited to the backsplice junction for probe binding, and one probe that could specifically target circH1. This would result in unobservable fluorescence. To mitigate this, we instead synthesized Cy5 labeled circH1 and transfected it in HeLa cells. By building a three-dimensional view using z-stacks, circH1 was found to be exclusively localized in the cytoplasm (Fig S6). A downside of this approach involved not studying the localization of native circH1 which could have a different landscape attributed to cellular modifications.

### circH1 is active *in cellulo*

Once we had established the *in vitro* activity of circM1, we next evaluated if these circular ribozymes are active inside the cells too. From a pool of pre-tRNAs^36^, 10 pre-tRNA were shortlisted for circH1 knockdown downstream effects. The pre-tRNAs were chosen to incorporate pre-tRNAs corresponding to a variety of amino acids, and both high and moderate expression profiles (Table S3). Since pre-tRNA already have low levels, we did not include pre-tRNAs with a low expression profile. Collectively, this ensured that the effects being studied were global and not limited to a narrow scope. Our initial hypothesis assumed circH1 to be catalytically active in HeLa cells and hence, knockdown of circH1 should lead to accumulation of pre-tRNAs as these pre-tRNAs cannot be cleaved into their mature isoforms (Fig. 6a). We employed RT-qPCR to monitor the levels of these pre-tRNAs using TBP as the housekeeping gene. The data from qPCR was analyzed using 2^−ΔΔCt^ method. Aligning with our hypothesis, 4 of these pre-tRNAs—A34, R4, Y8 and N14 were indeed upregulated up to 3 times their base levels (Fig. 6b). This is a direct evidence proving circH1 plays a role in the 5’ maturation of pre-tRNA. Since circH1 knockdown does not affect linH1 levels, the change in pre-tRNA levels is a direct result of circH1 downregulation. There are two possible pathways for circH1 participation in this maturation process—1. circH1 is catalytically active and is responsible for 5’ leader cleavage alongside linH1, 2. circH1 level has an inverse relation with pre-tRNA levels and acts as a signaling molecule for transcription of pre-tRNAs. If the latter is true, pre-tRNA levels would increase while the linM1 levels remain constant, contradicting the efficient use of molecular and energetic resources. Hence, it is more likely that circH1 is indeed catalytically active and cleaves the 5’ leader of pre-tRNA playing a direct and critical role in tRNA production.

**Figure 6.**
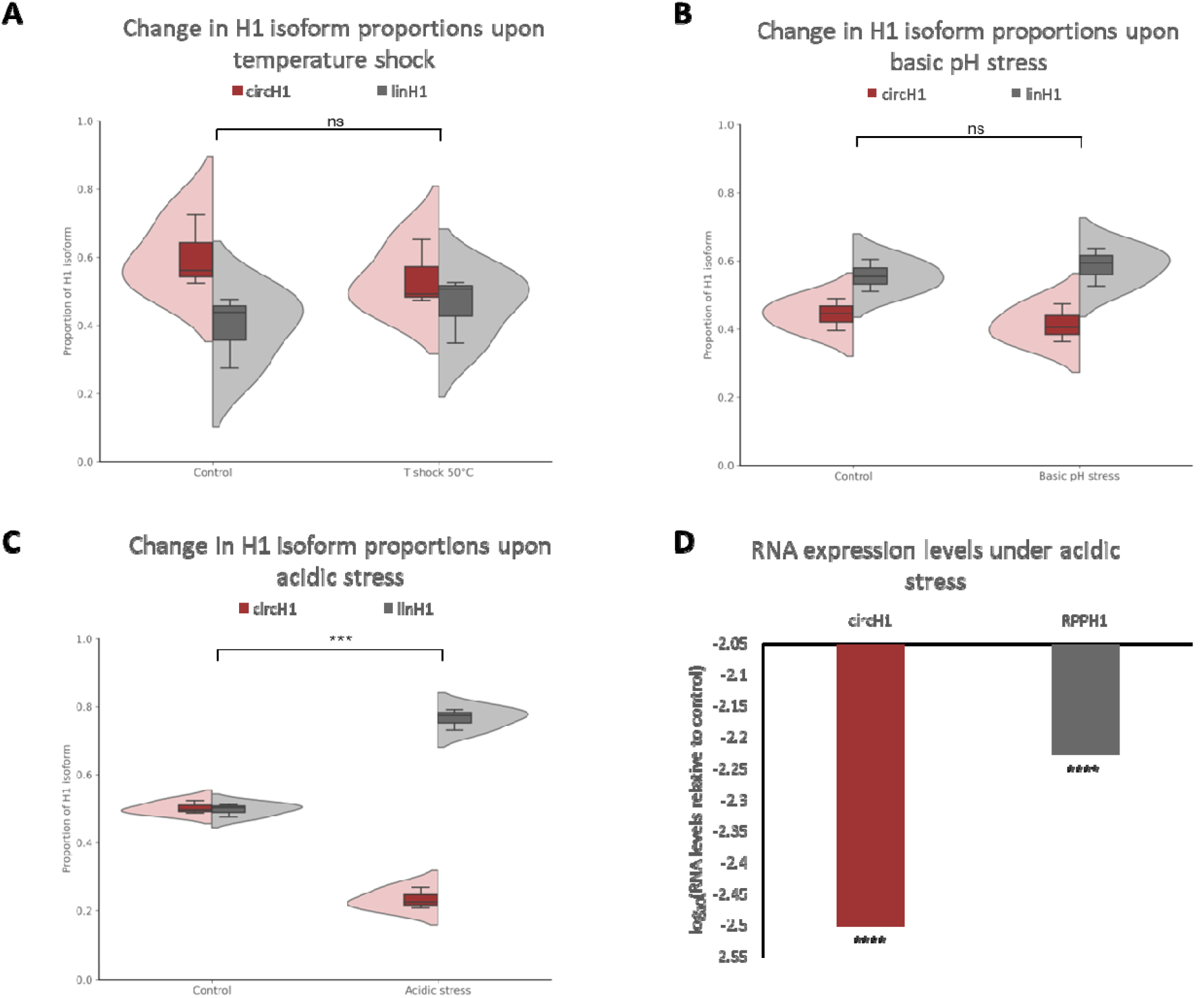
Acidic conditions suppress circH1 production. (A) No significant changes were observed betwee circH1:linH1 ratio upon treating HeLa cells to a temperature shock of 50°C for 24 hours. (B) No significant changes were observed between circH1:linH1 ratio upon treating HeLa cells to a basic pH shock of for 24 hours. (C) circH1:linH1 ratios were altered upon exposing HeLa cells to an acidic shock of 3.87 for 24 hours. (D) circH1 levels decreased more drastically than linH1 levels under acidic environment compared to control conditions. For quantitative analysis, n=3. Student’s t-test were performed to calculate p-values. p>0.05, ns; p≤0.05, *; p≤0.01,**; p≤0.001, ***.

Interestingly, one of the pre-tRNAs, U1 was downregulated by the same percentage of about 30% as circH1 upon siRNA knockdown (Fig. 6c). U1 corresponds to the amino acid, selenocysteine. U1 tRNA has a secondary structure different from all the other pre-tRNAs. Unlike other pre-tRNAs, U1 is first charged with serine followed by the phosphorylation of the hydroxyl group of serine by PSTK. SEPSECS then catalyzes the conversion to of this phosphorylated intermediate to selenocysteine^37^. Given the high correlation between circH1 knockdown and U1 pre-tRNA downregulation, the most likely mechanism involves circH1 acting as a regulatory RNA and signaling the slowing down of U1 transcription. Considering the two observed effects on pre-tRNA levels, circH1 possibly plays dual role of pre-tRNA maturation and transcription regulation.

### Acidic environment interferes with circH1 production

Once we had established the *in cellulo* activity in circH1, we then set forth to understand how the two H1 isoforms, circH1 and linH1, regulate under different cellular stresses. We exposed HeLa cells to three different stress conditions—acidic pH, basic pH and temperature shock, each for 24 hours. We then measured the transcript levels using RT-qPCR and analyzed the ratios by using M1 RNA as a spike-in control and 2^−ΔΔCt^ method.Under thermal and basic pH stress conditions, while the overall H1 transcript levels changed, the ratio of circH1 to linH1 remained relatively unchanged at about 0.5, suggesting an impact to transcription but not to circularization of H1 under these cellular stresses.

In contrast, acidic stress causes a significant shift in isoform abundance. Specifically, circH1:linH1 ratio decreased significantly from 1:1 to 1:3.3, indicating a preferential reduction of the circular isoform. Interestingly, while total levels of the parent RPPH1 transcript were markedly reduced, the decline in circH1 abundance was disproportionately greater than would be expected from a simple downregulation of transcription. This finding implies that acidic stress impairs the downstream circularization of H1 transcript, possibly by altering the activity, production or availability of factors necessary for circH1 biogenesis inside the cell.

## Discussion

Extensive research on RNase P in the past has revealed the importance of this enzyme in pre-tRNA maturation. A mitochondrial variant of RNase P RNA in human cell lines has been reported to perform the same function^38^ and protein-only RNase P enzyme was also discovered^39^. Herein, we have reported a circular variant of RNase P RNA in three different species. *In vitro* analysis of the bacterial variant, M1, showed superior performance of the circular isoform over the linear isoform under numerous conditions. While a deeper understanding of the mechanisms contributing to the difference in activities is needed, the closed loop structure is plausibly the primary contributing factor in enforcing the circRNA to fold in an active state under suboptimal conditions. Since we could not replicate the *in vitro* activity of H1 RNA using previously reported conditions (Fig. S4) and Rpr1 RNA has not been yet shown to be active *in vitro* in the absence of its cognate proteins, we did not investigate the *in vitro* activity of circH1 and circRpr1. This, however, does not negate the possibility of these circular variants being active *in vitro* and needs to be evaluated. Furthermore, we also found the two M1 isoforms perform differently in a biomarcomolecular rich serum, FBS. We hypothesize this inconsistency to be attributed to different molecular associations of the two isoforms in the serum. While the secondary structure of linear M1 is well established, to further understand the difference in the activity and behavior of the two isoforms, a structural analysis of the circM1 must be performed to substantiate findings in this study. *In vivo*, we found circH1 to be directly involved in pre-tRNA maturation in HeLa cells through BSJ-targeted siRNA knockdown of the circular RNA. Interestingly, circH1 also had a regulatory effect on the selenocysteine pre-tRNA. It must be noted that more experiments are needed to understand the exact pathway involved in the activity of circH1 in human cells and what other factors play a role in this mechanism. While we established circH1 is not produced through backsplicing, the mechanism of production of these circRNA in cells remains elusive. Firstly, evaluating the protein interactome can shed light on what proteins are involved in this ligation process. Protein knockdown can then be performed to examine the role of potential ligase candidates in circH1 synthesis. We note that the *in cellulo* activity of circM1 and circRpr1 remains unexplored as well. Well-designed systems regulating the levels of these circular RNA are required to study their downstream effect in more primitive biological systems. This study presents a novel and crucial role of circRNA across three organisms and opens avenues to exploring the presence of more circular ribozymes. We expect this study will lay the foundation for delving into the mechanisms behind the different behaviors of the linear and circular isoforms and a deeper understanding of the need for a circular variant.

## Conclusion

In this study, we have reported the first known circular ribozymes in three different species - *H. sapiens, E. coli* and *S. cerevisiae*. We found the bacterial variant, circM1, to perform superior to its linear counterpart, linM1, under various suboptimal conditions *in vitro*. We also established that these circRNAs are not relics of the past but possibly evolving mechanisms. Through sirRNA knockdown of circH1, we prove that this circRNA is not a result of backsplicing but a post-transcriptional process. Further, the siRNA knockdown also established the human variant, circH1, potentially has catalytic activity *in cellulo* and might be acting as a signaling pathway for selenocysteine pre-tRNA transcription regulation. As evident under basic conditions, circH1 facilitates linH1 in pre-tRNA maturation under stressful conditions. Collectively, these findings unveil a previously unknown and crucial function of circular RNA and opens avenues for a deeper understanding of RNase P dependent cellular pathways and diseases.

## Supporting information

Supplemental tables, list of oligomers used and additional images

