## Supplemental tables, list of oligomers used and additional images for "Unlocking efficiency: Native circular RNA surpass linear isoforms in RNase P activity": SI.docx

**pSupplemental Information**

**Table S1. List of chemicals and kits**

| **S. No.** | **Chemical/Kit** | **Manufacturer** |
| --- | --- | --- |
| 1. | Phusion High-Fidelity PCR Master Mix with HF Buffer | ThermoScientific |
| 2. | T4 RNA Ligase I | New England Biolabs |
| 3. | MEGAshortscript T7 Transcription Kit | Invitrogen |
| 4. | SequaGel UreaGel 29:1 Denaturing Gel System | National Diagnostics |
| 5. | 3M NaOAc solution pH 5.5 | Invitrogen |
| 6. | 5mg/mL Glycogen | Invitrogen |
| 7. | Ethanol | Pharmco |
| 8. | Guanosine Monophosphate | Sigma Aldrich |
| 9. | RNase R | Biosearch Technologies |
| 10. | Agarose | Sigma Aldrich |
| 11. | 2′-tBDSilyl Guanosine (n-PAC) CED phosphoramidite | Chemgenes |
| 12. | 2′-tBDSilyl Adenosine (n-PAC) CED phosphoramidite | Chemgenes |
| 13. | 2′-tBDSilyl Uridine (n-PAC) CED phosphoramidite | Chemgenes |
| 14. | 2′-tBDSilyl Cytidine (n-PAC) CED phosphoramidite | Chemgenes |
| 15. | 2'-tBDSilyl Cytidine (n-PAC) 3'-lcaa CPG 500Å | Chemgenes |
| 16. | Cyanine 5 Phosphoramidite | Glen Research |
| 17. | HPLC grade Acetonitrile | Sigma Aldrich |
| 18. | Trifluoroacetic acid | Fisher Chemicals |
| 19. | Triethyl amine | Thermo Scientific |
| 20. | Ammonium hydroxide | Sigma Aldrich |
| 21. | Methylamine | Sigma Aldrich |
| 22. | RNA purification cartridge | Glen Research |
| 23. | 2.0M Triethylamine acetate | Sigma Aldrich |
| 24. | RNA Quenching buffer | Glen Research |
| 25. | Dimethyl sulfoxide | Sigma Aldrich |
| 26. | DNA Clean & Concentrator-5 | Zymo Research |
| 27. | RNA Clean & Concentrator-25 | Zymo Research |
| 28. | Dulbecco’s Modified Eagle Medium (DMEM) | Gibco |
| 29. | Penicillin-Streptomycin (PenStrep) | Gibco |
| 30. | Fetal Bovine Serum (FBS) | Gibco |
| 31. | Luria-Bertani (LB) broth | Amresco |
| 32. | Quick-RNA Fungal/Bacterial MiniPrep | Zymo Research |
| 33. | Pierce™ Centrifuge Columns, 10 mL | Thermo Scientific |
| 34. | Millex®-GP Filter Unit 0.22 µm | Millipore Sigma |
| 35. | Triethylamine trihydrofluoride | Fisher Scientific |
| 36. | DBCO-Cy5 | AAT Bioquest |
| 37. | 10X PBS | Gibco |
| 38. | 0.25% Trypsin-EDTA (1X) | Gibco |
| 40. | Lipofectamine^TM^ 3000 | Invitrogen |
| 41. | Dextrose | Gibco BRL |
| 42. | Peptone | Becton, Dickinson and Company |
| 43. | Yeast extract | Becton, Dickinson and Company |
| 44. | SuperScript™ IV Reverse Transcriptase | Invitrogen |
| 45. | PowerTrack™ SYBR Green Master Mix for qPCR | Applied Biosytems |
| 46. | Formamide | Fisher Chemicals |
| 47. | Ethanol | Decon Labs, Inc |
| 48. | DAPI | ThermoScientific |
| 49. | CleanCap eGFP mRNA | Trilink Biotechnologies |
| 50. | Cyanine 5-Aminoallylcytidine-5'-Triphosphate (Cy5-CTP) | Trilink Biotechnologies |

**Table S2. List of instruments**

| **Name** | **Manufacturer** |
| --- | --- |
| Mermade 4 | LGC Biosearch Technologies |
| LSM 880 Confocal Microscope | Zeiss |
| Gel Imager | BioRad |
| Quant Studio 5 qPCR | Applied Biosystems |

**Table S3. List of DNA and RNA sequences**

| **Name** | **Sequence** | **Manufacturer** |
| --- | --- | --- |
| circH1_F | CCCATTATAGGGCGGAGGGA | IDT |
| circH1_R | TGGGAGTGGAGTGACAGGAC | IDT |
| circM1_F | GTCTGGTCAGCTTCAGGTGAA | IDT |
| circM1_R | TTCACCTGAAGCTGACCAGAC | IDT |
| circRpr1_F | GTCATTAGTGGCGAACCCGA | IDT |
| circRpr1_R | GGATCCTTCTGTAAACAGGTTTCT | IDT |
| M1_T7_F | TAATACGACTCACTATAGGAAGCTGACCAGACAGT | IDT |
| M1_R | AGGTGAAACTGACCGATAAGCC | IDT |
| circH1_340nt_R | CGGAGGAGAGTAGTCTGAATTGG | IDT |
| circM1_377nt_R | AGTCGCCGCTTCGTCGTCGT | IDT |
| circRpr1_369nt_F | CCAGCGTGGAACAGTGGTAA | IDT |
| circRpr1_369nt_R | AACAGCAGCAGTAATCGGT | IDT |
| pretRNA_Asp_F | TAATACGACTCACTATAGGACATGGTCCGG | IDT |
| pretRNA_Asp_R | TGGCGGTCCGGACGGGACTC | IDT |
| TBP_F | CACGAACCACGGCACTGATT | IDT |
| TBP_R | TTTTCTTGCTGCCAGTCTGGAC | IDT |
| Erse1_F | CGTTTGGTCAGGGCATCAAC | IDT |
| Erse1_R | ACTGAAGAGTGGCGTATGGC | IDT |
| 5.8SrRNA_F | CGGATCTCTTGGTTCTCGCA | IDT |
| 5.8SrRNA_R | GGAATACCAAGGGGCGCAAT | IDT |
| M1 template |  | TWIST Biosciences |
| Cy5-pATSerUG | GAUCUGAAUGGAGAGAGGGGGUUCAAAUCCCCCUCUCUCCGCCAC | In-house |
| pre-tRNAA34_F | GTCACCAAAGCGTTTTCCGA | IDT |
| pre-tRNAA34_R | CCGCTTCTTACTGGTGTCCC | IDT |
| pre-tRNAU1_F | TGCGCGGGAATAAGGTTGT | IDT |
| pre-tRNAU1_R | GGTTCGATAAGTAAGATTTAAGGCG | IDT |
| pre-tRNAR4_F | GCGTAAGGAGAGGGTGTGAG | IDT |
| pre-tRNAR4_R | CGCCCTCTCTTCCACCAAAA | IDT |
| pre-tRNAY8_F | CCTAAAGAATCCAAAACGAGGGT | IDT |
| pre-tRNAY8_R | CAGGCCCAGAAGAGACACTTT | IDT |
| pre-tRNAK9_F | CCACCCCGCCAATTACTAGA | IDT |
| pre-tRNAK9_R | TCCAGAACTCCAAGGTAGCA | IDT |
| pre-tRNAH11_F | CTATCATCGCTTGGCCTGTG | IDT |
| pre-tRNAH11_R | AAGTGGATAGCTGACGGCCT | IDT |
| pre-tRNAN14_F | CAGCCAAAATCGCCAACGC | IDT |
| pre-tRNAN14_R | CCAGGGTTCACAAAATGCGAC | IDT |
| pre-tRNAQ21_F | TTGGGTGAAGCGTACCAAGA | IDT |
| pre-tRNAQ21_R | CAAAAGGTCCAGGAAGAGGATA | IDT |
| pre-tRNAT10_F | TCCACTGAGAGATTCCACCG | IDT |
| pre-tRNAT10_R | AAGCCCTACCTCGGGTCTC | IDT |
| pre-tRNAI5_F | TCCTTTAAAACAGCAGCGCATT | IDT |
| pre-tRNAI5_R | TAAGTGAAGGGCCCCACTATG |  |

**Table S3. List of pre-tRNA shortlisted for investigation**

| **pretRNA** | **Readcount** | **Corresponding amino acid** |
| --- | --- | --- |
| K9 | 19926.29 | Lysine |
| N14 | 19062.82 | Asparagine |
| A34 | 13126.24 | Alanine |
| H11 | 11001.03 | Histidine |
| U1 | 10042 | Selenocysteine |
| Y8 | 9418.425 | Tyrosine |
| Q21 | 8496.88 | Glutamine |
| R4 | 7841.769 | Arginine |
| T10 | 7465.876 | Threonine |
| I5 | 3882.767 | Isoleucine |

**Methods**

All kits were used according to the manufacturer’s protocol unless otherwise mentioned.

*circRNA detection in different cell lines*

HeLa cells were grown in DMEM supplemented with 10% FBS and 1% PenStrep at 37°C in 5% CO_2_/95% humidity till 70% confluency. Total RNA was isolated using Quick-RNA MiniPrep.

DH5α strain of *E. coli* was grown in LB broth at 37°C. Cells were harvested in the log phase and total RNA was extracted using Quick-RNA Fungal/Bacterial MiniPrep kit.

W303 strain of *S. cerevisiae* was grown in 0.02 g/L dextrose, 0.02 g/L peptone and 0.01 g/L yeast extract at 30°C. Total RNA was extracted using Quick-RNA Fungal/Bacterial MiniPrep kit.

cDNA synthesis was performed using Superscript IV. Polymerase Chain Reaction was performed using divergent primers and Phusion High-Fidelity PCR Master Mix with HF Buffer and analyzed using 2% agarose gel.

*RNase R assay*

Based on the cell line, varying amount of total RNA was treated with RNase R in a 10 µL reaction in 1X RNase R reaction buffer. Control reactions were performed in the absence of RNase R enzyme. The reactions were precipitated using 0.3M NaOAc, 0.2 mg/mL glycerol and 3X volume of 100% Ethanol at -80°C overnight. The RNA was pelleted using centrifugation at 4°C at 20,600 rcf. RNA pellet was washed with 500 µL of 75% Ethanol by centrifuging at 14000 rcf at room temperature. The pellet was then air dried and resuspended in nuclease free water. cDNA synthesis was performed using Superscript IV and amplified by PCR with Phusion High-Fidelity PCR Master Mix with HF Buffer using convergent primers for linear controls and divergent primers for circular RNA. PCR products were analyzed using 2% agarose gels.

*Sanger sequencing*

Primers pairs were designed where one primer was specific to the backsplice junction and the other primer adjacent to it. cDNA from total RNA was used for PCR. The double-stranded DNA products were purified using DNA Clean & Concentrator-5 and sequenced through Eurofins Genomics and compared to sequences of the linear counterparts retrieved from RNA central.

*M1 RNA and 5´-Cy5-circH1 synthesis*

DNA templates corresponding to the required RNA sequences with T7 promoter sequence at the 5´ end were purchased from TWIST Biosciences, amplified using PCR and purified using DNA Clean & Concentrator-5. MEGAshortscript T7 Transcription Kit was used for RNA synthesis. 375 mM Guanosine Monophosphate (GMP) was added to the M1 synthetic reaction. This allowed the incorporation of monophosphate at the 5´ end. 0.5mM Cy5 labeled CTP was added to the reaction for synthesis of 5´-Cy5-circH1. The reaction was carried out at 37°C for 15 hours followed by DNase I treatment at 37°C for 15 minutes. The RNA was precipitated overnight, centrifuged, washed and air-dried as described above. M1 RNA was then analyzed using a 6% denaturing polyacrylamide gel.

*Synthesis of pATSerUG*

Model pre-tRNA, pATSerUG, was synthesized using solid-support oligonucleotide synthesis at 1 µM scale with DMT-ON using Mermade 4 synthesizer. The RNA was cleaved from the bead support and bases were deprotected in a single step using 1100 µL 1:1 ammonia/methylamine by shaking at room temperature for 4 hours. This solution was filtered using Pierce columns and the beads were washed twice with 300 µL water. Ammonia was evaporated under vacuum and the solution was lyophilized to obtain 2´ protected DMT-ON RNA. This was dissolved in 125 µL of DMSO and incubated in 60 µL of TEA and 75 µL of TEA/H_3_F to deprotect the 2´ hydroxyls. RNA was then purified and DMT was cleaved off using RNA purification cartridge. The obtained RNA solution was lyophilized and redissolved in nuclease free water. The obtained RNA was analyzed using 12% denaturing polyacrylamide gel.

*Synthesis of circM1*

M1 RNA was circularized using T4 RNA Ligase I and purified using a 6% denaturing polyacrylamide gel. The gel pieces were incubated in 0.3M NaOAc and 0.02 mg/mL glycogen and shaken overnight at 4°C. The solution was filtered through a 0.2-micron filter to remove the gel pieces and 3X volume of 100% ethanol was added to the filtrate. This was incubated at -80°C overnight, centrifuged and washed as described earlier.

*M1 activity assay*

0.24µM M1 RNA (linear or circular) M1 RNA was incubated in buffer (40mM MgCl2, 50mM Tris HCl pH 7.2, 5% PEG8000, 100mM NH4Cl), for 30 minutes at 37˚C followed by addition of 0.05µM pATSerUG pre-warmed to 37˚C. The reaction (2µL) was incubated at 37˚C for 40 minutes and quenched by addition of 8µL of 2X OrangeG in formamide. The solution was denatured at 90°C for 2 minutes and quickly cooled on ice for 1 minute followed by loading on a 20% denaturing polyacrylamide gel. Quantitation of gel bands was performed using Bio-rad ImageLab software. P-values were calculated using student’s t-test.

*Assay of circH1 in cellulo activity*

300k/well of HeLa cells were seeded in a 6-well plate in DMEM supplemented with 10% FBS and 1% PenStrep at 37°C in 5% CO_2_/95% humidity till 50% confluency. At 50% confluency, transfection of si-circH1 was performed. Old media was replaced with fresh DMEM. 100 µmol siRNA specific to circH1 was added to 125 µL of OptiMEM. The solution was vortexed and spun down, followed by a 15-minute incubation at room temperature. 0.4 µL of Lipofectamine RNAiMAX was added to 125 µL of OptiMEM and the solution was vortexed and gently spun down, followed by a 15-minute incubation at room temperature. The two solutions were mixed and incubated for 10 minutes at room temperature and then 250 µL of the prepared solution was eventually added to the well. After 48 hours, the media was removed, and cells were lysed using 300 µL of RNA Lysis Buffer and total RNA was extracted using Quick-RNA MiniPrep kit. The RNA was analyzed on a 2% agarose gel to determine its purity and intactness of the ribosomal bands.

1500 ng purified RNA was used as template for cDNA formation using SuperScript IV in 10 µL reaction. The synthesized cDNA was amplified using sequence specific primers and PowerTrack SYBR Green Master Mix for qPCR, employing TBP as the housekeeping gene. Samples were heated at 95°C for 10 minutes followed by 40 cycles of 95°C for 15 seconds and 60°C for 1-minute incubations. Samples were then incubated at 60°C for 1-minute and 95°C for 15 seconds. The data collected from qPCR was analyzed using 2^-ΔΔCt^ method and p-values were estimated using student’s t-test.

In cellular stress exposure experiments, HeLa cells were grown in 6-well plate as described above. Upon reaching 50% confluency, cells were exposed to cellular stress. For pH stress, the media was changed to DMEM supplemented with 10% FBS and 1% PenStrep, pH adjusted to either 3.87 (acidic pH stress) using 1N HCl, or (basic pH stress) using 1N NaOH and incubated at 37°C in 5% CO_2_/95% humidity for 24 hours, followed by total RNA extraction as described above. cDNA reactions were spiked with 0.5ng/µL with M1 RNA. RT-qPCR was performed as described above.

*Visualization of circH1 localization*

10K cell were seeded in one well of a 96-well plate. Upon reaching a 50% confluency, cells were transfected using 100ng Cy-5 circH1, eGFP and Lipofectamine MessengerMAX. circH1 and eGFP were added to 5µL of OptiMEM media, vortexed and briefly centrifuged. Similarly, another solution was prepared by mixing 0.3µL of the lipofectamine and 5µL of OptiMEM. Both solutions were incubated at room temperature for 15 minutes. The two solutions were then mixed, vortexed, briefly centrifuged and further incubated at room temperature for 10 minutes. Cells were replenished with fresh DMEM. The Lipofectamine-circH1-eGFP solution was then added to the cells while swirling the plate. After 24 hours, cells were fixed using 3.7% (vol./vol.) formaldehyde in 1X PBS and incubated at room temperature for 10 minutes. Then, the fixed cells were washed twice with 200 µL 1X PBS followed by permeabilization using 70% ethanol at 4°C for 1 hour. Following this, cells were stained with DAPI according to the manufacturer’s protocol.

HeLa cells were then visualized using confocal microscopy at 100X magnification using oil immersion in three channels—Cy5, eGFP and DAPI. Z-stacks were taken in an interval of 1µm and combined using ImageJ Fiji.

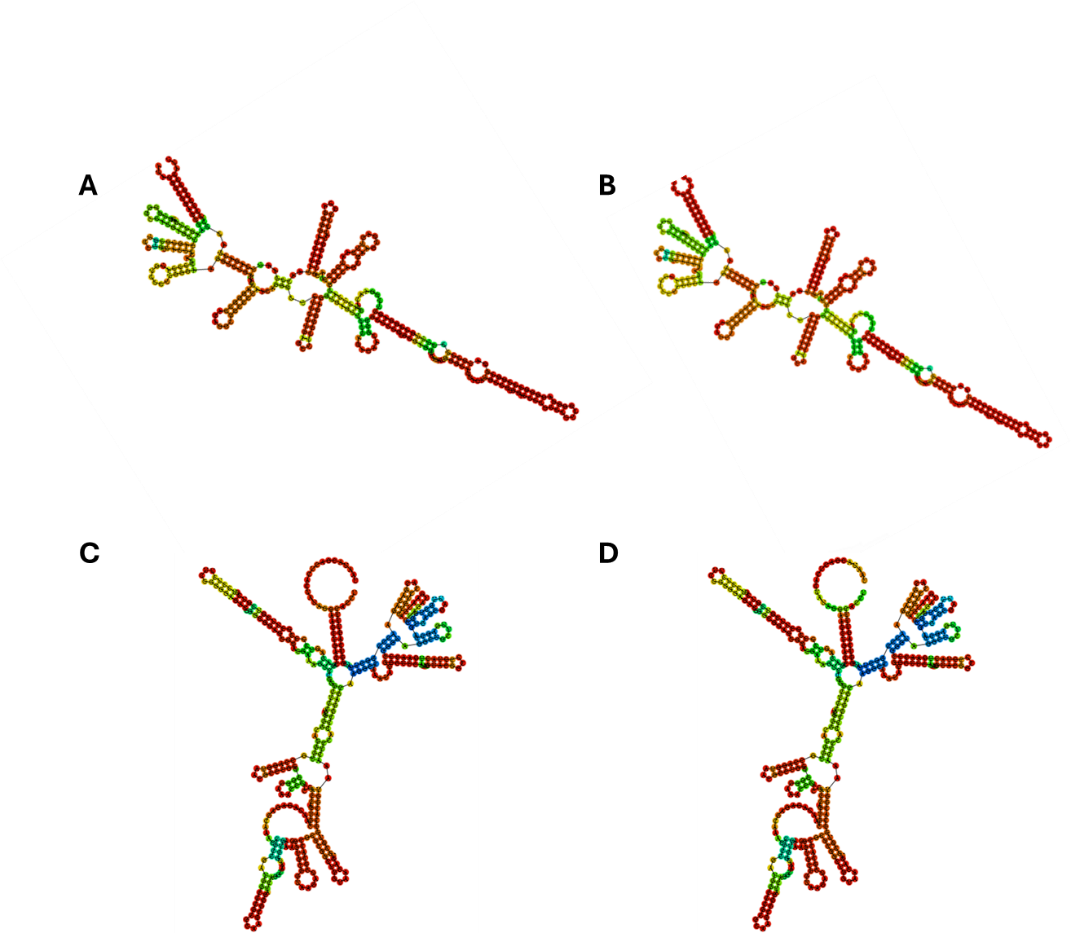

Figure S1. Secondary MFE structures of (A) linM1 and (B) circM1 and secondary centroid structures of (C) linM1 and (D) circM1 predicted using RNAfold

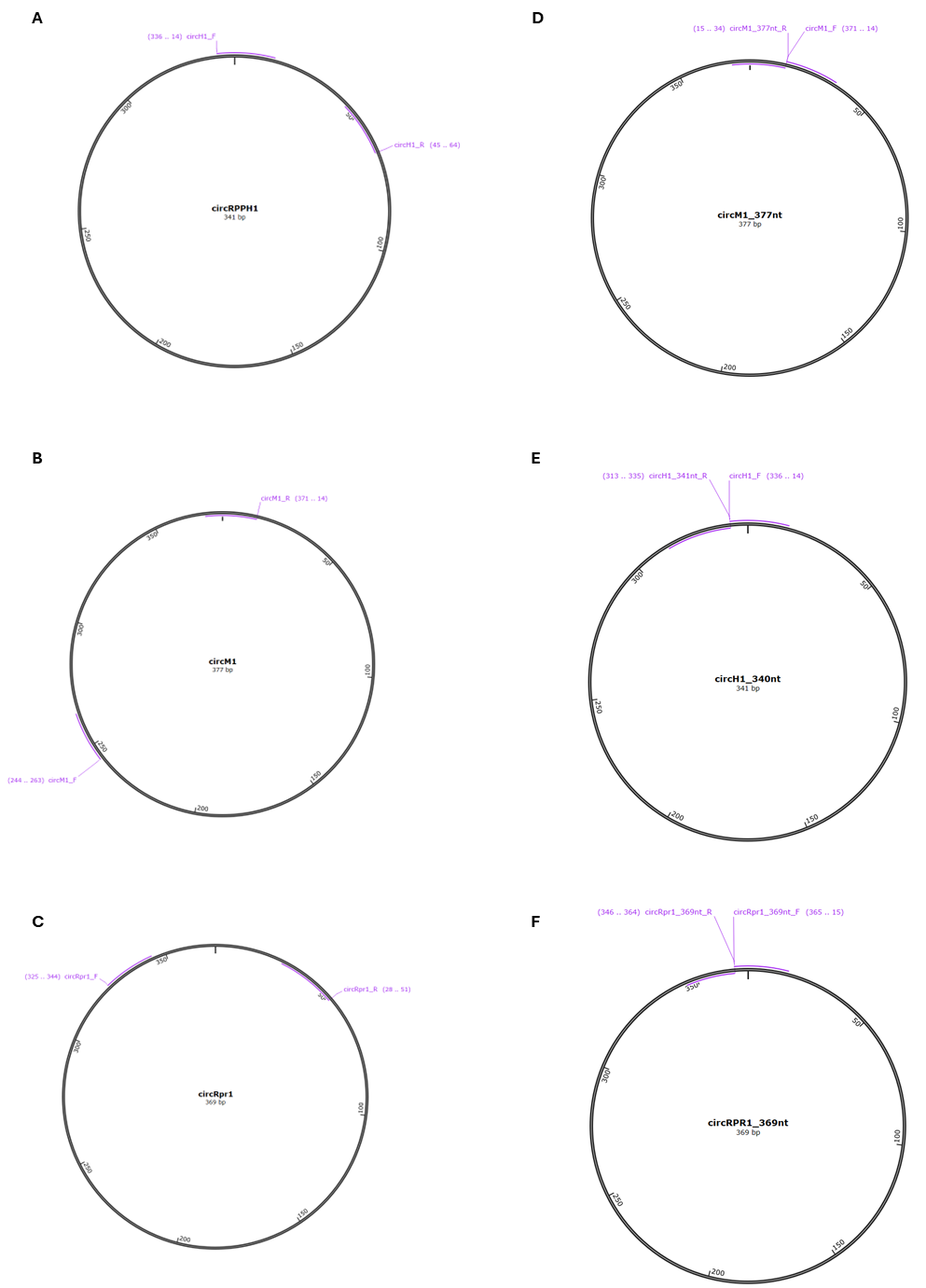

Figure S2. Positions of divergent primers used for detection of (A) circH1, (B) circM1 and (C) circRpr1. Positions of divergent primers used for cDNA synthesis followed by PCR amplification for Sanger sequencing of (A) circH1, (B) circM1 and (C) circRpr1.

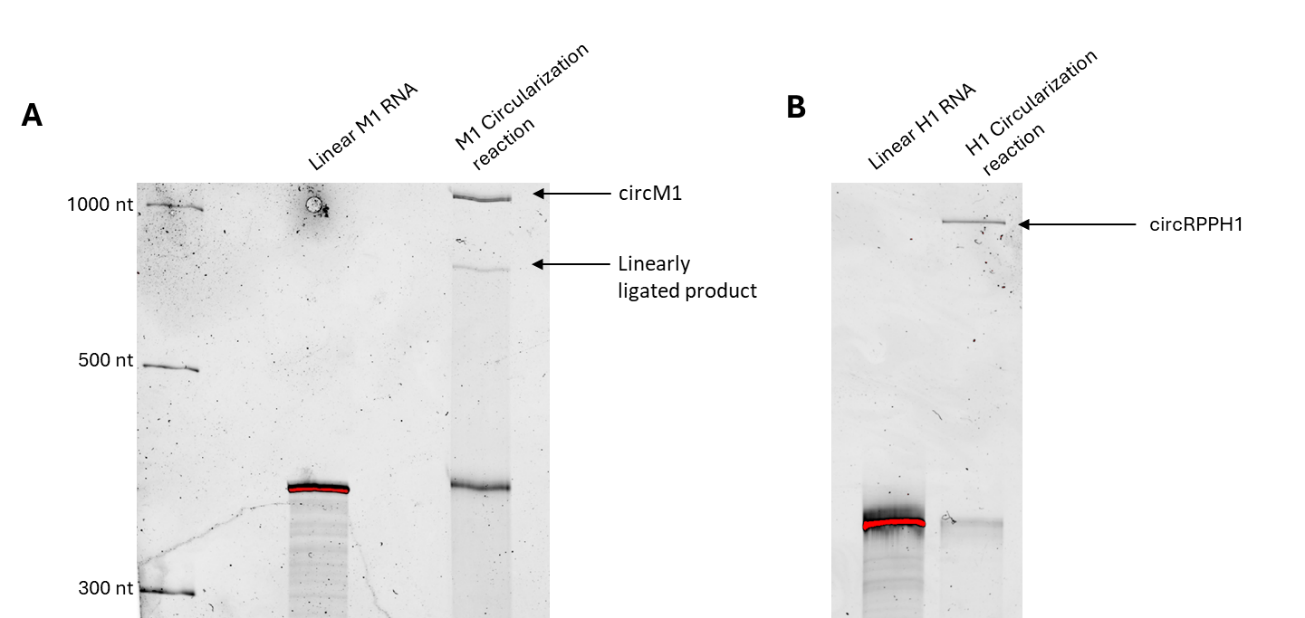

Figure S3. (A) circM1 and (B) circH1 were synthesized using T4 RNA Ligase I. Products were analyzed using 6% denaturing polyacrylamide gel.

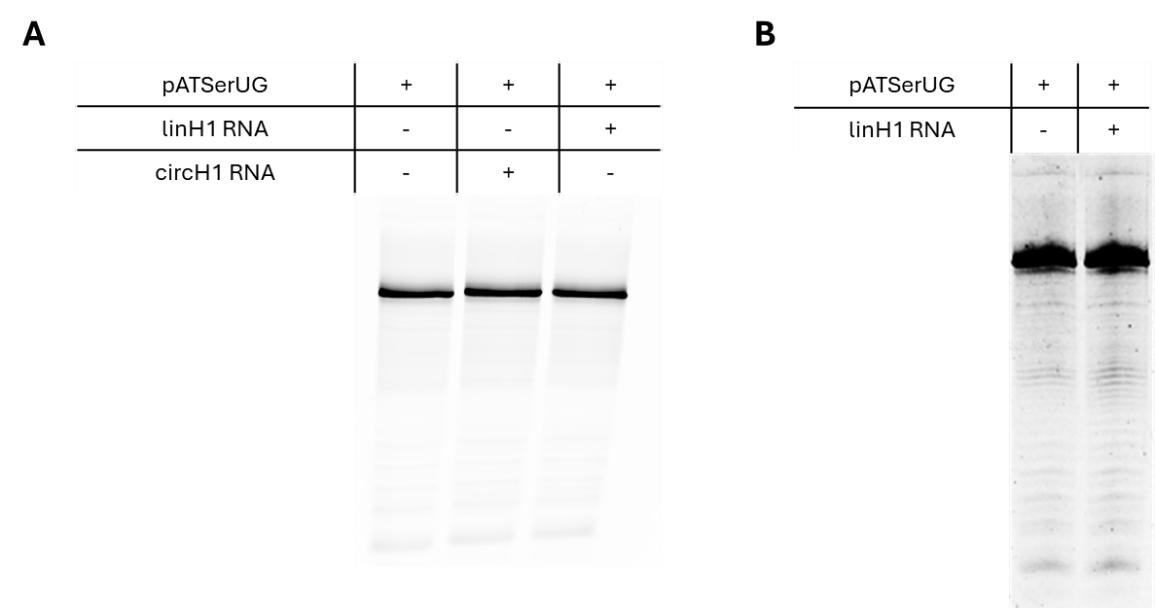

Figure S4. (A) Cleavage reaction of pATSerUG using linH1 and circH1. Buffer (40mM MgCl_2_, 50mM Tris HCl pH 7.2, 5% PEG8000, 100mM NH_4_Cl), 0.24µM M1 RNA, 0.05µM 5´Cy5-pATSerUG. M1 RNA was incubated in buffer for 30 minutes at 37˚C followed by addition of pATSerUG pre-warmed to 37˚C. *Reaction (2µL) was incubated at 37˚C for 40 minutes.* (B) Cleavage reaction of pATSerUG using previously published conditions.

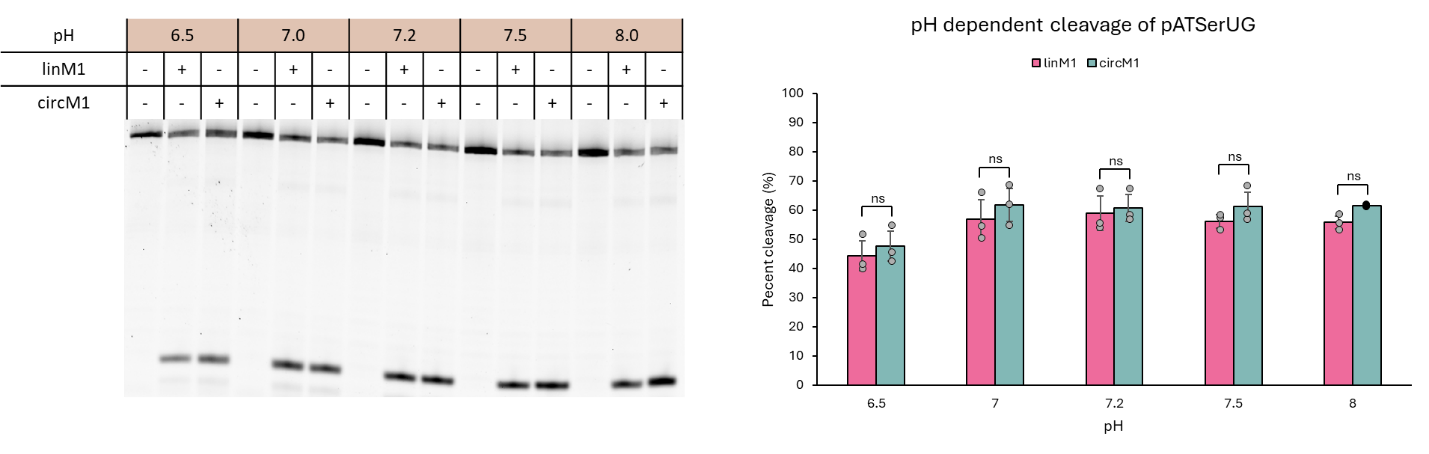

Figure S5. pH dependent cleavage of pATSerUG using linM1 or circM1. Buffer (40mM Mg^2+^, 50mM Tris HCl pH 6.5-8.0, 5% PEG8000, 100mM NH_4_Cl), 0.24µM M1 RNA, 0.05µM 5´Cy5-pATSerUG. M1 RNA was incubated in buffer for 30 minutes at 37˚C followed by addition of pATSerUG pre-warmed to 37˚C. Reaction (2µL) was incubated at 37˚C for 40 minutes.

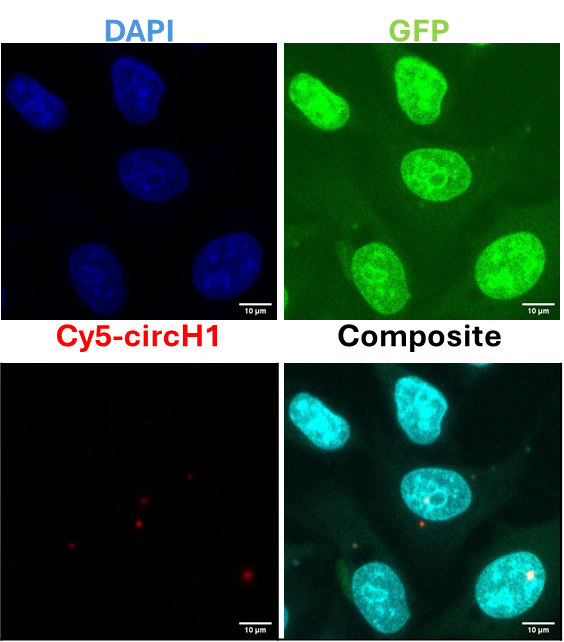

Figure S6. Cellular localization of transfected Cy5-circH1 in HeLa cells. 200ng of internally labeled Cy5-circH1 and 25ng of eGFP mRNA were transfected to HeLa cells grown in a 96-well plate at 40% confluency. After 24 hours, cells were fixed with formamide, permeabilized with ethanol and stained with DAPI. Images were taken at 100X magnification using an oil lens.

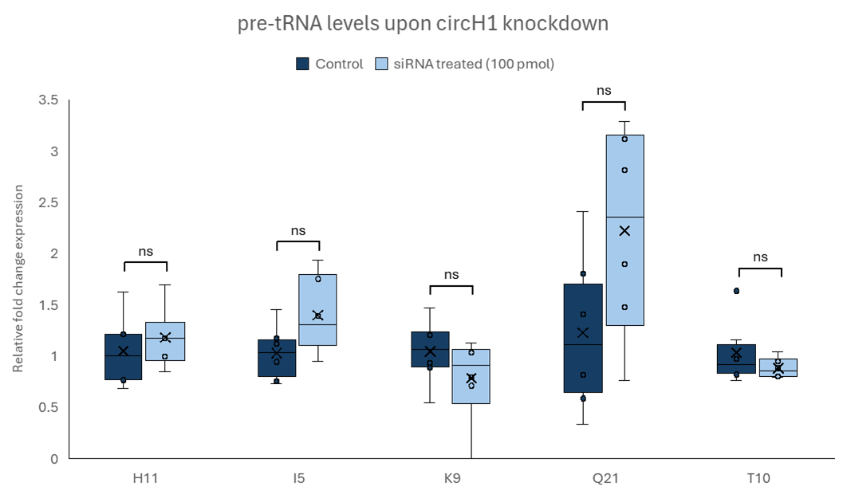

Figure S7. Some pre-tRNA levels remained unaltered by circH1 knockdown. For quantitative analysis, n=6. Student’s t-test were performed to calculate p-values. p>0.05, ns; p≤0.05, *; p≤0.01,**; p≤0.001, ***.
